# Analysing the structure of pathways and its influence on the interpretation of biomedical datasets

**DOI:** 10.1101/333492

**Authors:** Bram Burger, Luis Francisco Hernández Sánchez, Ragnhild Reehorst Lereim, Harald Barsnes, Marc Vaudel

## Abstract

Biochemical pathways are commonly used as a reference to conduct functional analysis on biomedical omics datasets, where experimental results are mapped to knowledgebases comprising known molecular interactions collected from the literature. Due to their central role, the content of the functional knowledgebases directly influences the outcome of pathway analyses. In this study, we investigate the structure of the current pathway knowledge, as exemplified by Reactome, discuss the consequences for biological interpretation, and outline possible improvements in the use of pathway knowledgebases. By providing a view of the underlying network structure, we aim to help pathway analysis users manage their expectations and better identify possible artefacts in the results.

## Introduction

In order to interpret the results of biomedical studies in a larger biological context, it is common to perform so-called pathway analysis. This can provide additional insight into the interactions between the detected compounds and possibly uncover underlying disease mechanisms (Garcia-Campos et al., 2015; Khatri et al., 2012). Pathways are defined as chains of biochemical reactions that together form high-level biological processes. The main participants of pathways are proteins, present in different states referred to as proteoforms (Smith and Kelleher, 2013; Regnier and Kim, 2017).

Our current knowledge of pathways and the molecular processes comprising them is consolidated in various knowledgebases as reviewed by Rigden et al. (Rigden et al., 2016), e.g. WikiPathways (Slenter et al., 2017), Ingenuity Pathway Analysis (qiagen.com), KEGG (Kanehisa et al., 2017), and Reactome (Milacic et al., 2012; Fabregat et al. 2018). Biases and knowledge gaps in pathway databases directly influence the results (Wadi et al., 2016), and insight into where our knowledge is lacking can be used to guide future research.

In this study, we systematically investigated existing pathway knowledge, with a focus on proteins and their interactions, using Reactome as a reference. The goal is to shed light on the structure and content of the data, and how this ought to influence the way pathway analysis is performed. For a comparison of the current pathway analysis approaches we refer the reader to (Garcia-Campos et al., 2015), as this is beyond the scope of this article.

Reactome is manually curated and contains detailed information on proteins (but also small molecules, RNA, DNA, carbohydrates, and lipids) connected to each other by chemical reactions, organized in a graph database that can be queried programmatically (Fabregat et al., 2017). Unless stated otherwise, our analysis should however be generic and it is anticipated that the findings also apply to other pathway databases.

Our results provide novel insight into the state of pathway knowledge, how it is structured, and indicate biases that may influence biomedical analyses. Finally, potential improvements in how bioinformatics tools interact with pathway databases are identified.

## Materials and Methods

Reactome was downloaded as graph database from reactome.org/download-data (version 58). Connecting to Neo4j (driver version 3.0.7) was done in Java 8 using the Neo4j Java Driver (version 1.0.6), and in R (version 3.4.2, (R Core Team, 2017)) *via* RNeo4j (White, 2016). Selections within the database were done by filtering EntityWithAccessionedSequence, Reaction, Pathway, and TopLevelPathway on speciesName: ‘Homo sapiens’, and ReferenceEntitiy on databaseName: ‘UniProt’. The networks presented in this manuscript can be created using PathwayMatcher (github.com/LuisFranciscoHS/PathwayMatcher). The code used to produce the results of this study can be found at github.com/bramburger/PathwayStructure.

Participation in reactions was selected by searching recursively for edges annotated with input, output, catalystActivity, regulator, regulatedBy, physicalEntity, hasMember, hasCandidate, and hasComponent. Direct participation of reactions in pathways was selected by searching for edges annotated with hasEvent and analogously, for indirect participation, although in that case the search was recursive. Gene Ontology terms were taken from the human complement of UniProt (The UniProt Consortium, 2017), downloaded on July 13th, 2017.

Radiality of each node was calculated by summing the reverse geodesic (i.e. one plus the diameter of the network minus the distance between the two nodes) for those nodes that can be reached, and subsequently dividing this sum by the diameter of the network times one minus the total number of nodes in the network. Integration was calculated in the same way but following the edges in reverse order (Valente and Foreman, 1998).

## Results

### Protein networks are becoming larger and denser

Maps of interactions between proteins are generally displayed in the form of protein-protein interaction (PPI) networks, where proteins are depicted as nodes and the interactions between them indicated by edges. Such graphs provide a model for biological knowledge at both local and global scales, i.e. they show the direct interactions between individual proteins as well as the overall network organization. A typical use case is to identify clusters of proteins specifically related to a given disease condition (Menche et al., 2015).

PPI networks can be constructed using experimental data, e.g. yeast two-hybrid (Y2H) screening or affinity purification coupled to mass spectrometry (AP-MS), or from interactions documented in the literature (Bork et al., 2004). The data contained in pathway databases fall in the latter category and are obtained either *via* manual curation or through automated literature mining, in contrast to databases of experimental inferred interactions, such as Biogrid (Oughtred et al., 2016) or Intact (Orchard et al., 2014).

By design, pathway networks differ from experimental PPI networks in three main aspects: (i) they have a higher granularity, i.e. they contain much more detailed information about the interactions, including, for example, information on the proteins’ post-translational state or sub-cellular localisation where the interaction occurs; (ii) they contain very different types of interaction, some of which are directed, including, for example, complex formation, catalysis, or inhibition; and (iii) given that the gathering of the data is dependent on manual curation, their growth is slower.

In **Figure 1**, we illustrate the growth pace by taking the Reactome database and locating the earliest literature reference for each protein interaction. The first documented reaction is from 1934, describing the interaction between two Haemoglobin subunits (Ferguson and Roughton, 1934). The network for 1934 thus contains only two nodes and two edges and is easily interpretable (**Figure 1A**). From 1960 to 2000 our knowledge about the network grew from 15 proteins in four separate components to a much larger network of 5,620 proteins across 70 components (**Figure 1B** to **1E**). In general, each year a substantially higher number of interactions was discovered compared to the number of proteins added to the network. Most of the interactions and proteins included were first mentioned in the literature between 1985 and 1995 (**Figure 2A**). The current network (**Figure 1F**) consists of 10,365 proteins (7,548 if excluding proteins not interacting with any other proteins), and over a million interactions, thus covering more than half of the human protein coding genes. The network has one densely connected main component, with slightly less well-connected periphery, and 85 smaller components (**Figure S1**).

**Figure 1:**
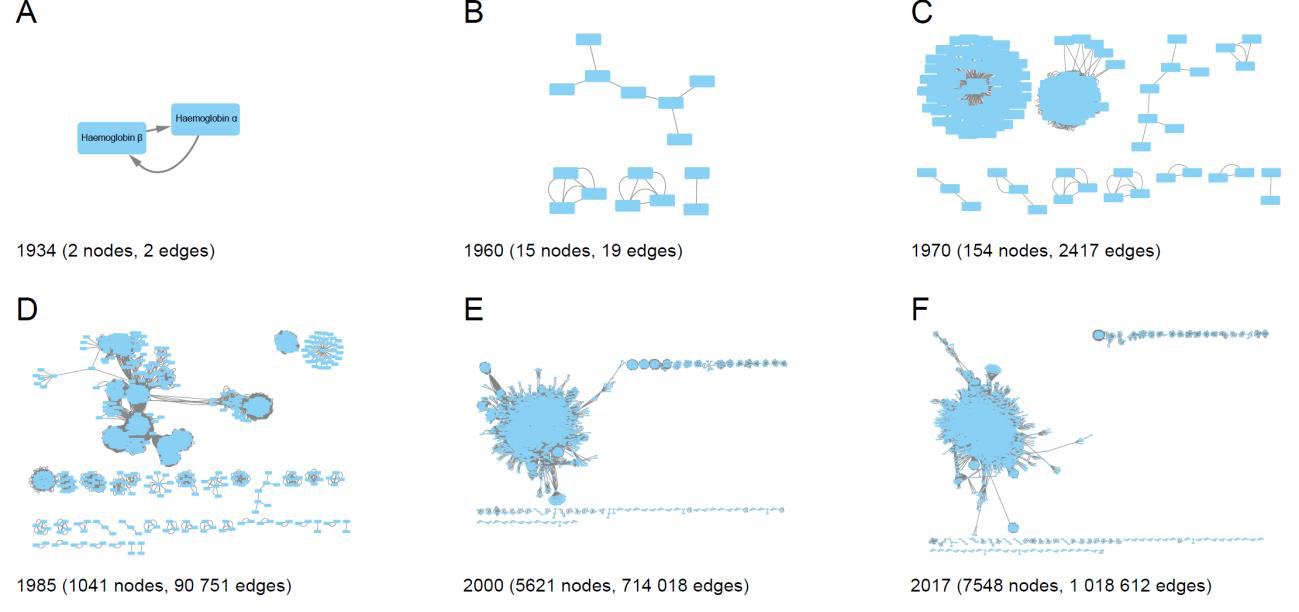
*Protein network evolution. For each year, only proteins participating in a reaction published in or before that year according to Reactome are included. A) 1934: only one reaction is documented, B) 1960: four connected components containing 15 proteins in total, C) 1970: larger, very dense components start to appear, D) 1985: several very densely connected components, connected to each other in various degrees, E) and F) 2000 and 2017: a single very densely connected component, with slightly less well-connected periphery and a number of smaller components. See main text for further details.*

**Figure 2:**
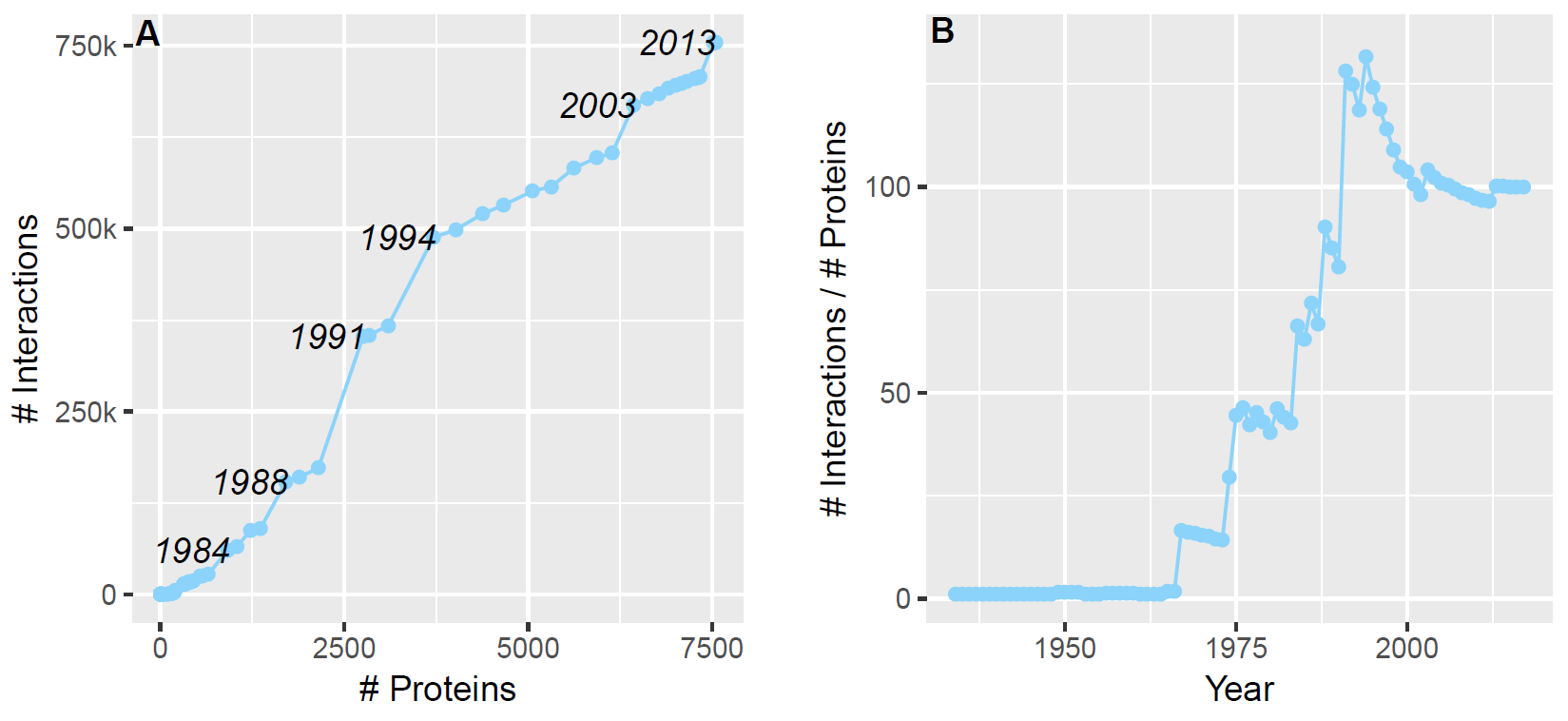
*Network size evolution. A) Number of proteins vs. number of interactions in Reactome. The number of interactions between two proteins and the number of proteins that interact with at least one other protein are plotted for each year, based on the earliest annotated publication. B) Average number of interactions per protein. The total number of interactions divided by the total number of proteins for each year, based on the earliest annotated publication.*

The average number of interactions per protein has steadily increased from 1965 onwards. However, after a sharp drop between 1995 and 2000, the average number of interactions per protein seems to have stabilised around 100 (**Figure 2B**). There are also large differences in the number of annotated interactions per protein. e.g. the maximum number of interactions for a single protein is 1,329, while 367 proteins have only one documented interaction.

The increase in the number of annotated proteins and interactions has made inference concerning the affected processes in the biological system more challenging, i.e. the gain of additional data has come at the price of increased complexity.

### From protein interactions to pathways

Reactome defines interactions as chemical reactions (e.g. binding reactions, transportation or modifications) which have an input and an output, and possibly a catalyst and/or regulator. In the following, we define two proteins as interacting when they appear together in a reaction and where one of the proteins is the output of the reaction. The other protein is either an input, a catalyst, or a regulator. In this model, all interactions are therefore directed. A binding reaction, where two (or more) proteins form a complex (**Figure 3A**), is defined as having the pair of proteins both as input and output, producing two edges between the proteins, one in each direction (**Figure 3B**).

**Figure 3:**
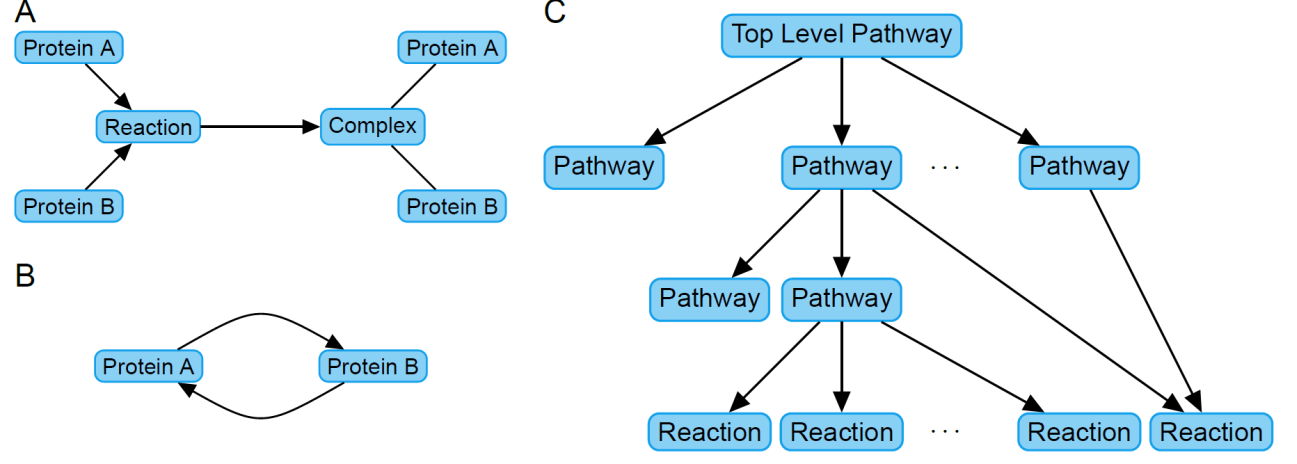
*Protein interactions in Reactome. A). Proteins can simultaneously be input and output of a reaction, e.g. a binding reaction producing a complex. B) In the resulting networks, proteins in a complex will have an edge between them in both directions. C) Pathway hierarchy generic structure: reactions compose pathways, pathways may also group together more specific pathways, and top-level pathways group related pathways to the more generic processes.*

Complexes are important in the interpretation of interactions, as all proteins in a complex need to be present in a given folding and modification state to form a functional complex, and subsequently, a complex can only perform its task(s) when all relevant proteins are bound together. A related concept is entity set, where one of a set of candidate proteins has to be present to perform a given task, i.e. proteins in entity sets are thus interchangeable (Alberts, 1984; Fabregat et al., 2017).

A protein that only acts as the input of reactions will only have outgoing edges and can be thought of as a starting point for a biological process, while a protein that never acts as the input of a reaction will be an end point of a process. Chains of reactions do not necessarily follow one after the other. For example, when a protein-complex is the input of a process, the complex first has to be generated, which can be a separate process.

Reactions having specific functions when chained together are grouped into pathways. A pathway is thus a description of a process, with the corresponding reactions making the process happen. This can be a very specific process, consisting of only a few reactions, or general pathways described as collections of more specific pathways, with potentially additional reactions connecting them. The pathways are then grouped together in even more general pathways, where pathways that do not belong to any other pathway are referred as top-level pathways (**Figure 3C**).

In total, Reactome contains 2,051 pathways, with 1,400 of these not having any sub-pathways, i.e. being the most specific pathways consisting only of reactions. 654 pathways contain at least one other pathway and most of these contain only a few other pathways (**Figure S2**). However, the top-level pathways, and some of their sub-pathways, have numerous pathways nested inside them. The largest top-level pathway is Signal Transduction (2,444 proteins), while the smallest is the Circadian Clock (21 proteins) (**Figure 4A**). Thus, finding ten differentially quantified proteins in the Circadian Clock pathway is clearly not as likely as finding the same number of proteins in the Signal Transduction pathway. This is why the size of the pathways is a key parameter when analysing and comparing pathways.

**Figure 4:**
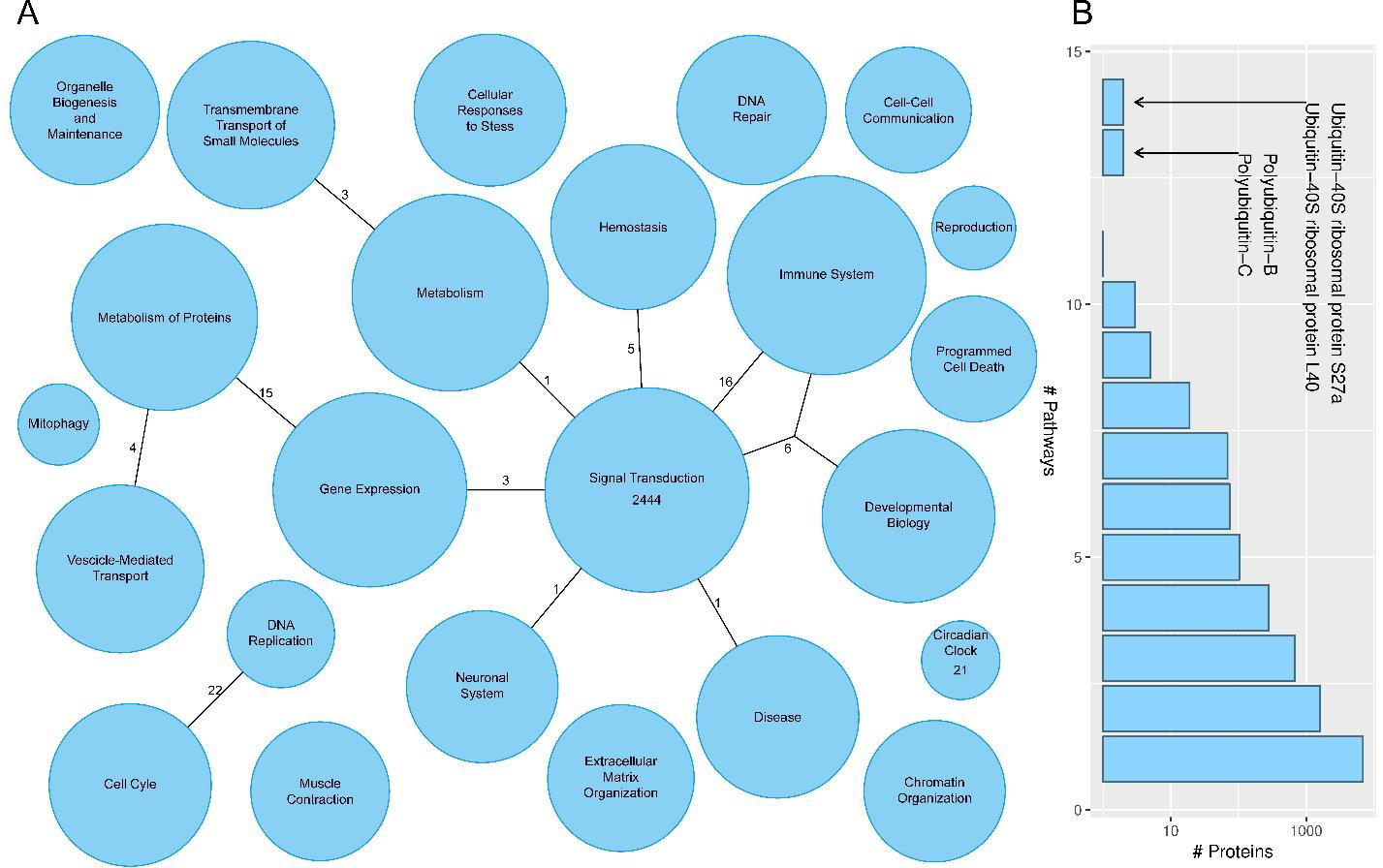
*Pathway sizes. A) Sizes of top-level pathways and the number of shared pathways. The size of the circles is proportional (log scale) to the number of proteins in each top-level pathway. The largest and smallest pathways have the number of proteins annotated. Edges between pathways indicate the number of sub-level pathways they have in common. B) Distribution of the number of top-level pathways a protein participates in. The y-axis is in log-scale. Proteins participating in more than half of the top-level pathways are annotated.*

While it is intuitive that a pathway can have multiple sub-pathways (as biological processes often consist of multiple, more or less distinct, sub-processes), the opposite is also possible, i.e. sub-pathways being part of multiple parent pathways. This can be interpreted as processes being re-used by several distinct processes. However, this is not very common, with only seven pathways being sub-pathways of more than two other pathways, and 41 having exactly two parent pathways. The pathways that descend from more than two other pathways are listed in **Table S1**. Identifying components of such ubiquitous pathways does not allow identifying the higher-level biological mechanism at play, leaving ambiguity in the pathway inference.

The nested structure is important when interpreting pathway analysis output, as high-level pathways may have easily understandable descriptions but they are most often not very specific. On the other hand, for very specific pathways the other pathways it belongs to have to be taken into account when interpreting their place in the biological system of interest. When multiple pathways are found it becomes important to compare the different pathways to each other and see whether (and how) they are connected.

1,390 pathways directly contain at least one protein, with 271 of them also containing a sub-pathway, while 429 pathways only contain proteins indirectly *via* their sub-pathways, thus not containing any proteins directly. However, most pathways containing at least one protein only contain a few proteins (**Figure S3A**). While there are only eight pathways directly containing more than 200 proteins, the largest pathway directly contains close to 500 proteins.

It is also possible to include the number of proteins in nested pathways when counting how many proteins are part of a given pathway (**Figure S3B**). This way, a pathway may artificially appear of interest due to the contribution of different sub-pathways. For example, identifying two proteins that participate in the Immune System pathway, but occur in completely different parts of the network, is less meaningful than proteins clustering in a specific region of the immune system.

Just as a pathway can be part of several other pathways, a protein can participate in multiple pathways. At the most general level, more than half of the proteins in Reactome participate in only one top-level pathway (**Figure 4B**) and can be of high interest due to their specificity. Note that none of the proteins participate in all 24 top-level pathways, i.e. there are no proteins which have a role in all general processes. However, four proteins (ubiquitins) participate in more than half of all top-level pathways.

An indication of how many processes a protein is potentially involved in is obtained by counting all pathways a protein participates in (including sub-pathways of the given pathway). This way, most of the annotated proteins participate in less than a dozen pathways (**Figure S3C**), and few (618) participate in more than 25 pathways. The four ubiquitin proteins participating in more than half of the top-level pathways are the only proteins to participate in more than 150 pathways.

One can also count only the pathways a protein directly participates in. This indicates how many specific processes a protein is involved in, while each specific process can itself occur in multiple more general processes. This way, most proteins (8,680) participate in less than six pathways (**Figure S3D**), with the four ubiquitin proteins being again the only proteins participating in upwards of 100 pathways. While the proteins participating in few pathways may have more specific functions, it should be noted that the proteins participating in many pathways can also potentially affect separate pathways in different equally unique ways.

When a protein participates in multiple pathways, it may interact with different sets of proteins depending on the function it is performing. The blocked diagonal of the adjacency matrices, shown in **Figure 5A** and **B**, indicates that there are both dense and less dense clusters of interacting proteins in the pathways, while the differences in blockiness of the pathways indicate that some pathways have denser clusters of interacting proteins than others.

**Figure 5:**
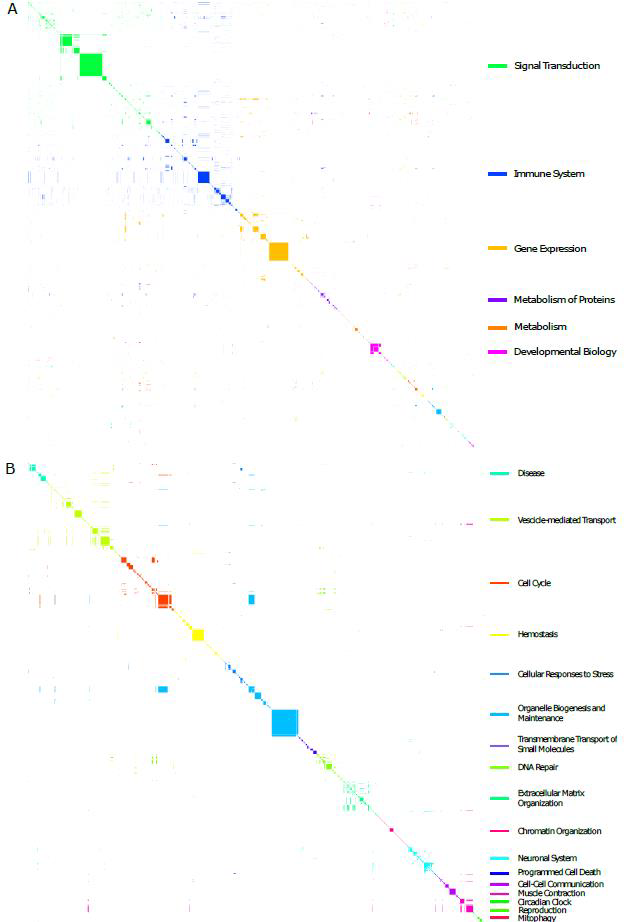
*Adjacency matrix showing the edges between protein interactions and pathways. Each row/column is associated with a specific protein; a dot indicates an interaction between two proteins. Interactions are colored by the largest top-level pathway they participate in. A) the complete adjacency matrix. B) only proteins not participating in one of the six largest pathways. Note that adjacency matrices may give a slightly biased view due to coloring by the largest pathway. Reactions/interactions can happen in multiple pathways.*

### A global view on protein interactions

Out of the 10,212 proteins in Reactome, 7,548 interact with other proteins. In addition, 4,539 of those can form complexes with other proteins. Most do so with only a few other proteins. Looking at interactions in general, most proteins have many interactions with other proteins, e.g. there are numerous proteins (4,795) with over 50 interactions, and quite a few (56) with more than a thousand interactions (**Figure S4**). Each of the four ubiquitin proteins mentioned above can have more than 2,000 interactions. In this jumble, it is therefore of great importance to filter out the interactions that are relevant for each specific study.

By looking at the current network (**Figure 1F**) it is possible to distinguish different types of nodes with the naked eye, based on their connectivity to other nodes. To better define these types, the radiality and integration of each node was calculated, where radiality provides an indication on how easily a node can reach other nodes, and integration gives an indication as to how easily other nodes can reach it (Valente and Foreman, 1998). The range for both metrics are 0 to 1, with higher values for radiality indicating that a protein can reach more proteins through less reactions.

Our results show that proteins can roughly be divided into four categories (**Figure 6**): (i) isolated proteins: both radiality and integration near zero (399 proteins); (ii) main component proteins: relatively high values for both radiality and integration (5,998 proteins); (iii) start of chain proteins: low integration, non-low radiality (385 proteins); and (iv) end of chain proteins: low radiality, non-low integration (766 proteins). As expected from **Figure 1** and illustrated in **Figure 6**, the radiality and integration of proteins has increased over time as the network was growing, and the prevalence of isolated proteins has diminished.

**Figure 6:**
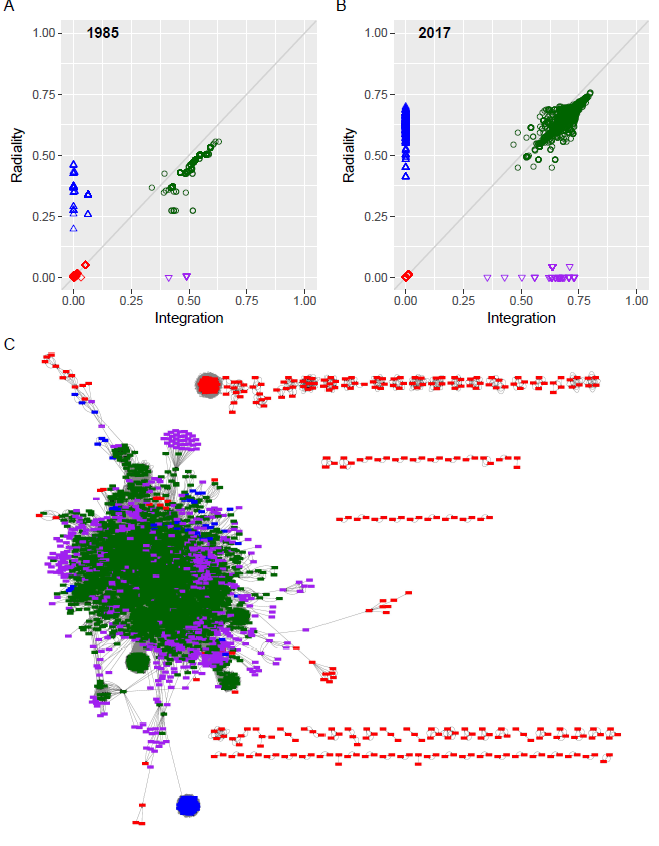
*Radiality and integration of the proteins in the network of 1985 (A) and 2017 (B, C). The PPI network for 1985 uses only interactions which earliest annotated literature reference is in or before 1985. Red diamonds: isolated proteins; green circles: proteins in the main cluster; blue upwards-pointing triangles: start of chain proteins; purple downwards-pointing triangles: end of chain proteins; grey diagonal: the line where radiality equals integration. (C) displays the entire network with nodes colored according to the categories identified in (B).*

Isolated proteins are part of small components not connected to the main component and thus can only reach a very small number of other proteins, or are in the periphery of the main component, but have few interactions (**Figure 6C**). The start and end chain proteins are at the periphery of the main component and are either at the beginning or at end of chains of reactions, respectively. Proteins that form the start of chains are able to reach many other proteins, but due to the directionality of the interactions cannot in turn be reached by many other proteins. Similarly, proteins that are at the end of chains are able to be reached by many other proteins but cannot themselves reach many other proteins.

The first important distinction is between the isolated proteins and the rest. When the proteins of interest are all isolated proteins, it will probably not make much sense to rely on network analyses to uncover biological context. Further interactions can then be obtained from experimental PPIs, and further investigation may be required to identify the functions and interactors of these proteins. Another potentially important distinction is between the start-of-chain, end-of-chain and main-component proteins. Deregulation or function impairment of the proteins might have different implications or likely causes depending on which type of protein they are.

## Discussion

During the recent decades our understanding of proteins and their interactions has steadily increased, and there are continuous ongoing efforts to curate and annotate this knowledge into publicly available knowledgebases. It is therefore reasonable to assume that with increasing knowledge it has become easier to view study specific findings in a larger biological context. On the one hand this is true, there are more proteins that we *can* put in a biological context. But on the other hand, the network has grown so large and dense that it has become increasingly challenging to identify the relevant interactions and processes.

The network seems to evolve towards a denser network, potentially consisting of a single large and closely connected component. In such a network it becomes vital to look at the interactions at two levels: the local interactions and the global context. Local interactions are the direct connections between the proteins, constituting the basic steps of the biological processes, essential for our understanding of how these processes work. At the same time, it is important to take into account where in the network the interactions take place. For example, when a protein at the start of a chain of reactions is affected by a disease, there may be less possibilities for the biological system to compensate. Conversely, when a protein somewhere in the middle is affected, there may be more opportunities to find alternate ways of achieving the given biological function. Finding redundancies in the system around affected proteins or proteins with homologous functions can possibly allow uncovering biological mechanisms more specifically and identify druggable targets.

Our findings also indicate large differences between pathways. This may be explained by some processes being more complex than others and thus requiring a greater number of proteins and steps to be executed and regulated. However, some processes have also received a greater amount of scientific attention, and the more research is carried out in a specific field the more detailed its annotation. The degree to which these two elements affect the differences in the annotation of pathways is uncertain, but it is clear that they ought to be taken into account when analysing and comparing pathways.

Rolland et al. found that the dense zones in the protein map may be the result of biases in the curation process, and not due to biological properties (Rolland et al., 2014). The curation seems to focus mainly highly and widely expressed proteins, which tend to already be well-connected in knowledgebases. Such biases can be managed through additional experiments and come as an important reminder that biological networks are tools for navigation and not a substitute for experimental validation.

Interestingly, this strategy of extending the network is encouraged by the Reactome curator guidelines (wiki.reactome.org/index.php?title=New_Reactome_Curator_Guide), and such a bias has an impact on what can be learned using pathway analysis, meaning that well-studied proteins will most likely only be put in the most well-known contexts. Similarly, there will be a greater number of annotations for proteins playing (potential) roles in well-researched diseases, and there can thus potentially be a bias towards the mechanisms involved (Schaefer et al., 2015). However, our comparison of the gene ontology (The Gene Ontology Consortium, 2016) annotations of the proteins in Reactome against those in UniProt revealed no obvious biases (**Figure S5A** and **B**).

One major simplification in the current pathway analyses is the use of a gene-centric model, where the proteoforms originating from the same gene, are modelled as a single node. Proteoforms can be due to, for example, amino acid variation, splice variants, sequence processing and folding, and post-translational modifications, potentially influencing a protein’s function and interaction partners. While the theoretical number of possible proteoforms is orders of magnitude larger than the number of experimentally observed proteoforms, the number of biologically relevant proteoforms is still unknown (Aebersold et al., 2018). Reducing all potential proteoforms to a single node in the protein network greatly simplifies biological knowledge, and is likely to reduce our ability to model highly specific protein-protein interactions.

As an example, the “RAC-alpha serine/threonine-protein kinase” protein (UniProt identifier P31749) is annotated to participate in 29 pathways in Reactome. If one knows that the protein is not modified, it is only annotated to participate in 10 pathways. Furthermore, if the protein has a modified tyrosine at coordinate 315 or 326 there is only one annotated pathway to inspect. This simple example indicates that asking more specific questions can provide dramatically more specific results. However, this is not always the case, e.g. when the same protein has a modified threonine at coordinate 308 the number of possibly affected pathways is still as high as 26.

When looking only at the interactions between pairs of proteins, important information is lost regarding participation in protein complexes and entity sets. Detecting only a single protein from a complex may indicate that the protein was not used in the specific reaction, although this conclusion is less likely to be correct when the other proteins in the complex are harder to detect. Given that many reactions depend on the interaction between protein complexes and/or entity sets, it is therefore essential to look beyond the interactions between pairs of proteins when doing pathway analysis. For example, when a protein interacts with a set of proteins, which in turn interacts with another protein, the dynamics are completely different when the set of proteins in the middle is a complex versus an entity set.

One should also keep in mind that proteins not annotated in a pathway database cannot be analysed in the context of pathways. Currently Reactome contains 10,212 human proteins, only just above half the number of human coding genes. Given this rather large gap, there seems to be a potential bias towards proteins that are ‘easier to find’, i.e. proteins having more annotation and/or better evidence for their existence (**Figure S5C and D**).

Finally, it is important to underline that even if a set of proteins is not annotated as interacting in a pathway database, their interaction may yet be known, but lacks proven involvement in one of the biological processes reflected in the collection of pathways – or this information was not curated yet. The process of choosing proteins and pathways for curation could be randomised to make it less biased, giving all molecules the same chance to get annotated in the pathways, but that would come at an increased cost for curation teams, as new expertise is needed for each new biological area to review.

Being able to understand project specific findings in a larger biological context is a key goal in the biomedical sciences, and with the increasing amount of knowledge available in public pathway knowledgebases this objective is increasingly achievable. However, our results indicate that the growing complexity of the protein network and its structural biases present major challenges for this field of research. Through better understanding of the pathway network structure, correction for biases in analyses, and improvement of the models for complex biological systems, we are confident that the accuracy of pathway analyses will constantly improve, providing biomedical scientists with ever expanding understanding of complex biological systems.

## Acknowledgements

BB, RRL, and HB are supported by the Bergen Research Foundation. HB is also supported by the Research Council of Norway.

## Author Contributions

Conceptualization: HB, MV; Methodology BB, LFHS, RRL, HB, MV; Software BB, LFHS; Writing – Original Draft BB, HB; Writing – Review & Editing BB, LFHS, RRL, HB, MV; Visualization BB, HB; Supervision HB, MV; Project Administration HB, MV; Funding Acquisition HB.

